# A generic risk assessment model for animal disease entry through wildlife: The example of highly pathogenic avian influenza and African swine fever in The Netherlands

**DOI:** 10.1101/2022.04.25.489353

**Authors:** Michel J. Counotte, Ronald Petie, Ed G. M. van Klink, Clazien J. de Vos

## Abstract

Animal diseases can enter countries or regions through movements of infected wildlife. A generic risk model would allow to quantify the risk of entry via this introduction route for different diseases and wildlife species, despite the vast variety in both, and help policy-makers to make informed decisions. Here, we propose such a generic risk assessment model and illustrate its application by assessing the risk of entry of African swine fever (ASF) through wild boar and highly pathogenic avian influenza (HPAI) through wild birds for the Netherlands between 2014-2021. We used disease outbreak data and abstracted movement patterns to populate a stochastic risk model. We found that the entry risk of HPAI fluctuated between the years with a peak in 2021. In that year, we estimated the number of infected birds to reach the Dutch border by wild bird migration at 273 (95% uncertainty interval: 254-290). The probability that ASF outbreaks that occurred between 2014 and 2021, reached the Dutch border through wild boar movement was very low throughout the whole period; only the upper confidence bound indicated a small entry risk. On a yearly scale, the predicted entry risk for HPAI correlated well with the number of observed outbreaks. In conclusion, we present a generic and flexible framework to assess the entry risk of disease through wildlife. The model allows rapid and transparent estimation of the entry risk for diverse diseases and wildlife species. The modular structure of the model allows to add nuance and complexity, when required or when more data becomes available.

## Introduction

Emerging and re-emerging animal diseases that affect livestock can be introduced into a country or region through movements initiated by humans, such as movement of live animals, products and people (Simons et al., 2019). However, many of these diseases can occur in wildlife populations as well, and thus movement of wildlife can act as possible entry route (Smith et al., 2017). This movement is often complex and based on many external factors that vary between regions and time such as availability of feed (Morelle et al., 2015), variation in temperature (Klinner & Schmaljohann, 2020) and climate change (Visser, Perdeck, van Balen, & Both, 2009).

Wildlife has played an important role in the transmission of several recent emerging and re-emerging animal diseases. For example, introductions of highly pathogenic avian influenza (HPAI) into the Netherlands were mainly attributed to migrating wild birds (Engelsma, Heutink, Harders, Germeraad, & Beerens, 2022). Seasonal migration patterns seem to drive the introductions of this virus (Xu, Gong, Wielstra, & Si, 2016). For instance, the 2005 spread of HPAI from Russia to the Black sea basin followed the spatiotemporal pattern of duck migration from Siberia (Gilbert et al., 2006), which was confirmed by phylogenetic analyses (Kilpatrick et al., 2006). In 2020, the HPAI outbreak in The Netherlands was preceded by outbreaks of the same virus strain on breeding grounds in Kazakhstan and Russia, leading to the conclusion that autumn migration of waterfowl likely caused the introductions (Beerens et al., 2021).

Similarly, for the entry of African swine fever (ASF) in European countries, wildlife plays an important role. Dispersal of wild boars is considered the highest risk for ASF introduction and spread in Europe (De la Torre et al., 2015; de la Torre et al., 2022). The number of ASF outbreaks in Europe has been increasing over the last decade (European Food Safety Authority et al., 2020; Linden et al., 2019), as has the number of affected countries. Where ASF virus spread by wild boar is mostly slow (approximately 50 km/year) (Bosch et al., 2017), long-distance jumps have resulted in outbreaks in wild boar in the Czech Republic in 2017, Belgium in 2018, and Italy in 2022, suggesting a human-mediated introduction rather than introduction by actively migrating wild boar (Sauter-Louis et al., 2022). Movement of wild boar has proven difficult to manage and the persistence and reinfection through carcasses have plays an important role in the continued transmission of ASF in Europe (Cukor et al., 2020; Woźniakowski, Pejsak, & Jabłoński, 2021).

Assessing the risk of introduction is a first step in the risk assessment that helps prioritize and plan preventive measures to mitigate the entry risk of emerging animal diseases, and surveillance activities for early detection. Risk assessment provides an ‘objective and defensible method’ of assessing the risk that pathogens pose to a country or region (World Organisation for Animal Health, 2021). Generic risk assessment tools allow for the rapid and transparent comparison of the risk across multiple diseases. The number of pathways addressed by these tools, however, varies largely and not all tools have incorporated wildlife movements as a pathway for disease incursion (de Vos et al., 2020). Most of these risk assessments are qualitative (EFSA, 2017) or semi-quantitative (Condoleo et al., 2021; Kyyrö, Sahlström, & Lyytikäinen, 2017; Roberts, Carbon, Hartley, & Sabirovic, 2011; Roelandt, Van der Stede, D’Hondt, & Koenen, 2017) and some of them rely heavily on expert knowledge. Simons et al. (2019) and Taylor et al., (2020) tried for a more quantitative approach where the wildlife risk was based on density and habitat suitability raster maps (Simons et al., 2019; Taylor et al., 2020). The latter assumes that the movement of wildlife is more likely towards raster cells where the habitat is more suitable. This approach was used to assess the incursion risk of ASF and might be appropriate for wild boar movement (Taylor et al., 2020). However, bird migration is hard to capture in a similar way, since their flight paths span over larger distances, and there are large seasonal fluctuations in their behaviour. This means that the generalizability of those models is limited, and detailed wildlife abundance data is required, which is often not available. Disease-specific risk models to assess the entry risk of HPAI via wild bird migration are mostly spatially explicit. For example, Kosmider et al (2016) assessed the entry risk of HPAI in the United Kingdom (Kosmider et al., 2016) using a risk score approach informed by the overlap of wild bird and poultry abundance, producing a risk map. Similarly, Martinez et al. (2011) used a spatial approach to identify areas of high risk, to perform risk-based surveillance (Martinez et al., 2011). These models tend to give a more detailed insight in the spatial distribution of risk, but at the expense of generalizability to other diseases.

In contrast to bespoke models, a generic risk model would allow to quantify the entry risk via wildlife movements for different diseases and wildlife species, despite the vast variety in both. Here, we describe a generic model to assess the entry risk of animal disease through wildlife and demonstrate its workings with the example of the entry of HPAI through wild birds and ASF through wild boar for the Netherlands.

## Materials and methods

In the risk assessment model described below, we estimate the risk of “entry” of a pathogen into a new territory through wildlife movements. We define the entry risk as the expected number of infected animals that reach the border of a territory per unit of time.

We first describe the generic model: The modelling approach, and the model structure and calculations. We then assess the model by assessing the entry risk of HPAI through wild birds and ASF through wild boar for the Netherlands. For the two diseases, we describe the input data, parameterization, model assumptions, and the validation of results.

All analyses were performed in R version 4.0.5 (R Core Team, 2021).

### Modelling approach

We considered that disease outbreaks that occur beyond the borders of a land or region could pose an entry risk through wildlife movements. This risk depends on geographic proximity and properties of the disease, such as duration of infectiousness, and the behaviour of the animals carrying the disease. We abstracted these complex disease and behaviour properties into a simple, generic set of variables: ‘directionality’, ‘distance’, ‘duration’ and ‘abundance’ to define different ‘behaviour groups’, accommodating for different movement patterns ranging from, for example, home range to migratory behaviour. Multiple ‘behaviour groups’ combined allowed us to mimic more complicated movement patterns. ‘Directionality’ is the probability that the outbreak spreads in a certain cardinal or intercardinal direction, the directions of an 8-wind compass rose. This can be considered as the direction the infected animal or group of animals decides to travel. ‘Distance’ is the total distance an infection can travel via wildlife. This is not limited to a single animal or group of animals; it also allows for unobserved transmission chains in the wildlife population. The model assumes that infected animals move in a single direction and travel in a straight line; the direction and distance may be different though in each iteration based on sampled values. ‘Duration’ is the period (in days) over which the transmission chain or infection can persist. ‘Abundance’ represents the monthly number of animals in each behaviour group; this parameter allows us to take into account fluctuations in abundance, for example, due to seasonal patterns. We defined the seasonal/temporal abundance within a behaviour group (monthly/temporal abundance denoted by *T*_*abundance*_) relative to the month with the highest abundance (See parameterized example).

Abundance was also used to define the relative probability of behaviour groups by month based on their relative contribution to the combined abundance of all behaviour groups (behaviour group abundance denoted by *B*_*abundance*_; See parameterized example below). This, together with the seasonal fluctuations of animal numbers over time (*T*_*abundance*_) allows to capture ‘seasonality’ or temporal changes over the year. Behaviour groups can represent different species, different age classes, or other properties that require a distinction into different behaviour groups.

### Model structure and calculations

The model considered the entry risk of every reported disease outbreak outside the area for which the risk assessment is performed. First, from the geolocated outbreak, we calculated the distance to the closest border point of the area of interest (The Netherlands in the parameterized example) and the heading in degrees for the shortest straight line from the outbreak to that border point (Figure 1) using the *geosphere* package (Hijmans, 2019). The direction was then reduced to one of eight cardinal or intercardinal directions.

**Figure 1.**
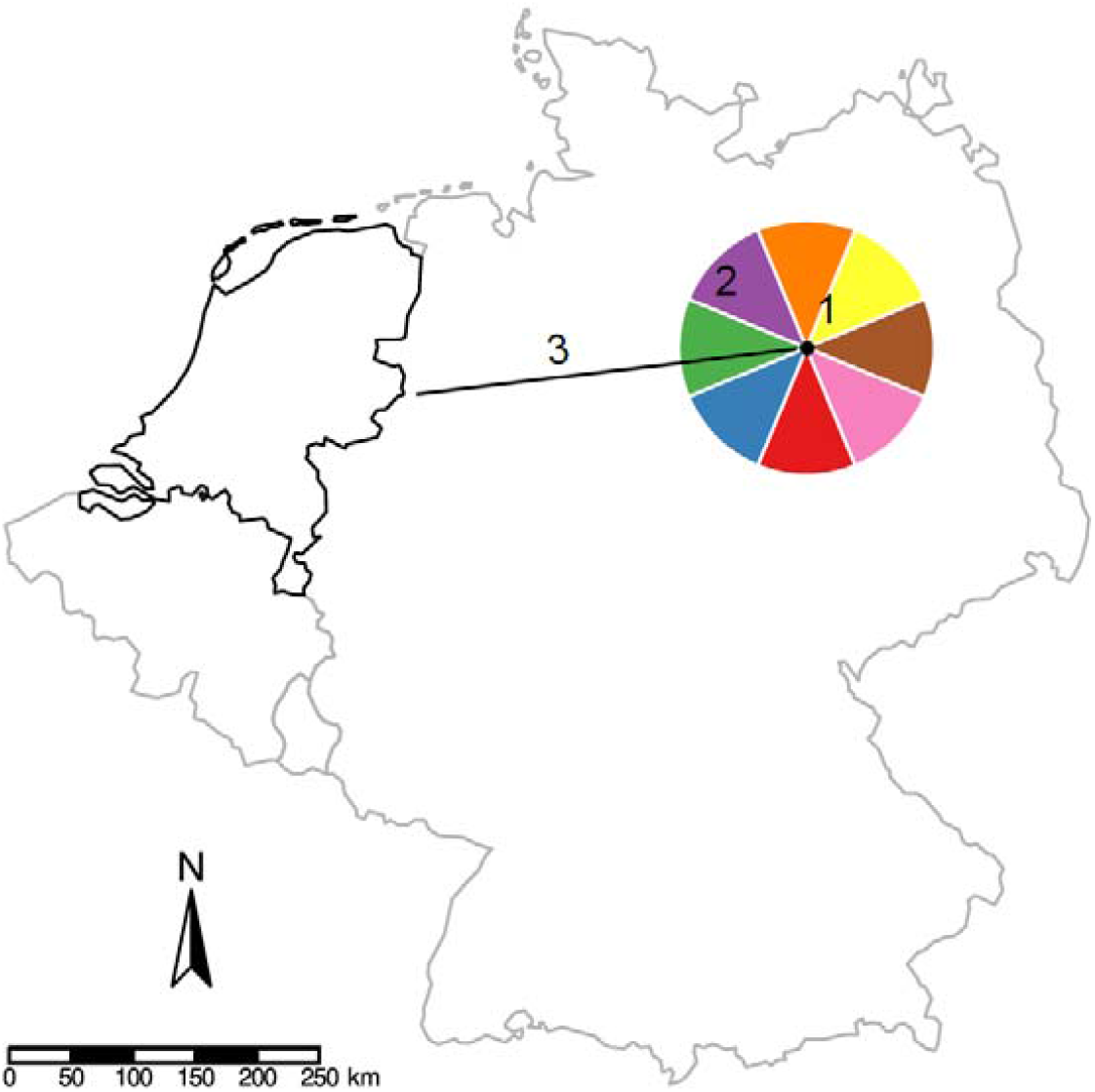
Model structure. An example for a hypothetical outbreak in March in Germany. From the hypothetical outbreak (1), the direction (2) and the shortest distance to the border (3) of the Netherlands were calculated. In this case, the direction was West and the distance was 325 km. Then, the behaviour was sampled based on the probability distribution of March. From the sampled behaviour, distance and direction were sampled. Only when subsequently a westward direction and an effective travel distance larger than 325 km are drawn, the outbreak is considered to have reached the Netherlands.

Second, one of the behaviour groups was sampled using the relative probability/abundance of behaviour groups of the month in which the outbreak is observed (*B*_*abundance*_). The behaviour group then provided the probability distribution of the effective *distance*, the probability distribution of the cardinal and intercardinal *direction*, and fixed values for the *duration* and the temporal abundance (*T*_*abundance*_). Third, the distance and direction were sampled from these distributions. If the sampled direction corresponded with the direction in which the closest border point is located and the sampled distance was larger than the distance to that point, we considered that the outbreak reached the border. Last, we calculated the daily entry risk. The entry risk (*R*_*i,t*_) by outbreak (*i*) and time (*t*) is the product of the outbreak size, i.e. the number of infected animals reported in the outbreak (*size*) and the monthly relative abundance (*T*_*abundance*_) within the sampled behaviour group, divided by the duration (*duration*) in days over which the transmission can take place (equation 1).

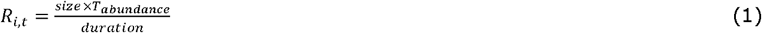

Thus, we considered the outbreak to pose an entry risk from the observation date until the end of the infectious duration. The total daily risk (*R*_*overall* _*t*_) of all *n* outbreaks is the sum of the risk of each outbreak per day (equation 2).

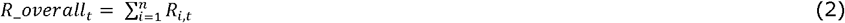

To calculate the monthly or yearly risk, we summed the daily risk over the number of days per month or year. Because of the stochastic nature of the model, we ran 100 iterations of the model and collected the median and 95% uncertainty interval of the entry risk.

From the model, we collected the following outputs: 1) The number of infected animals reaching the border of the Netherlands per month and year. 2) The individual contributions of a) ‘Behaviour groups’, and b) source countries to the monthly and annual entry risk.

### Model input data

We retrieved data on disease outbreaks from the Food and Agriculture Organization’s (FAO) Emergency Prevention System (EMPRES-I) (available from: https://empres-i.apps.fao.org/). In this database, disease outbreaks and a minimal set of characteristics are indexed by the FAO. For the indexed outbreaks, the geolocation (latitude, longitude), the observation and reporting date, and the size of the outbreak and affected species are reported. We considered outbreaks that were labelled as ‘wild’ in the species description to have occurred in wildlife. The database provides outbreak data starting from 2005. We extracted all reported ASF and HPAI outbreaks between January 1, 2014 and December 31, 2021. We opted for this period, since in the period 2005-2014, few HPAI outbreaks were reported in the Netherlands. Similarly, during that period ASF was mainly circulating in Russia, and introductions in the European Union (Poland) only started in 2014. As described above, for all disease outbreaks, we calculated the distance to the border of the Netherlands, and the cardinal or intercardinal direction. In the baseline model, we considered both outbreaks that were reported in wildlife as well as outbreaks reported in domestic animals. Furthermore, we considered all outbreaks to be of the same size (size = 1), due to a large proportion of outbreaks with missing information on the number of infected animals.

### Infection and animal behaviour parameters

#### Highly pathogenic avian influenza

To model the entry risk of HPAI through wild birds, we defined two behaviour groups: 1) a short range movement that occurs year round without a preferential direction, and 2) seasonal migration where an influx of birds into the Netherlands peaks in autumn (Figure 2D, and Supplement 1 for additional details). For the first, we assumed that infections through wild birds can pose a risk for 15 days (duration), and can travel 30 kilometres per day (assumed to be normally distributed with a standard deviation (sd) of 5 km), resulting in a maximum median distance of 450 km (*distance*, Figure 2B). For the second, the migrating birds, we considered 13 bird species of the Anatidae family to be relevant for the introduction of HPAI in the Netherlands, i.e. the Greater White-fronted Goose (*Anser albifrons*), Eurasian Wigeon (*Anas penelope*), Gadwall (*Anas strepera*), Common Teal (*Anas crecca*), Mallard (*Anas platyrhynchos*), Northern Pintail (*Anas acuta*), Garganey (*Anas querquedula*), Northern Shoveler (*Anas clypeata*), Red-crested Pochard (*Netta rufina*), Common Pochard (*Aythya ferina*), Tufted Duck (*Aythya fuligula*), Common Coot (*Fulica atra*), and the Brant Goose (*Branta bernicla*) (Velkers et al., 2021). Except for the Garganey, that winters in Africa, the majority of the birds originate from breeding grounds in a North, Northeast and East direction from the Netherlands (*directionality*, Figure 2C, based on the Migration Mapping tool (https://euring.org/research/migration-mapping-tool) and expert input). Their flight range can span up to 5000 km (assumed to be normally distributed with a sd=750 km) (*distance*) and we assumed that they can pose a risk over 25 days (*duration*), taking into account the persistence of infection within a transmission chain. In Supplement 1, we provide additional details on the selected bird species and their behaviours that led to the abstraction described above. The number of migratory birds changes due to seasonal migration patterns which is reflected in the temporal abundance (*T*_*abundance*_) (Figure 2D). The October migratory peak population was taken as reference value (100%) to calculate the temporal abundance of migratory birds for other months. Based on expert input, we assumed that the abundance of the ‘short range’ birds was constant at 5% of the October migratory peak population. The relative abundances of both behaviour groups were used to set the relative probability of the behaviours over time (Figure 2A). The relative probability of the behaviour groups (*B*_*abundance*_) was thus calculated as the relative abundance of each behaviour group divided by the relative abundance of both behaviour groups.

**Figure 2.**
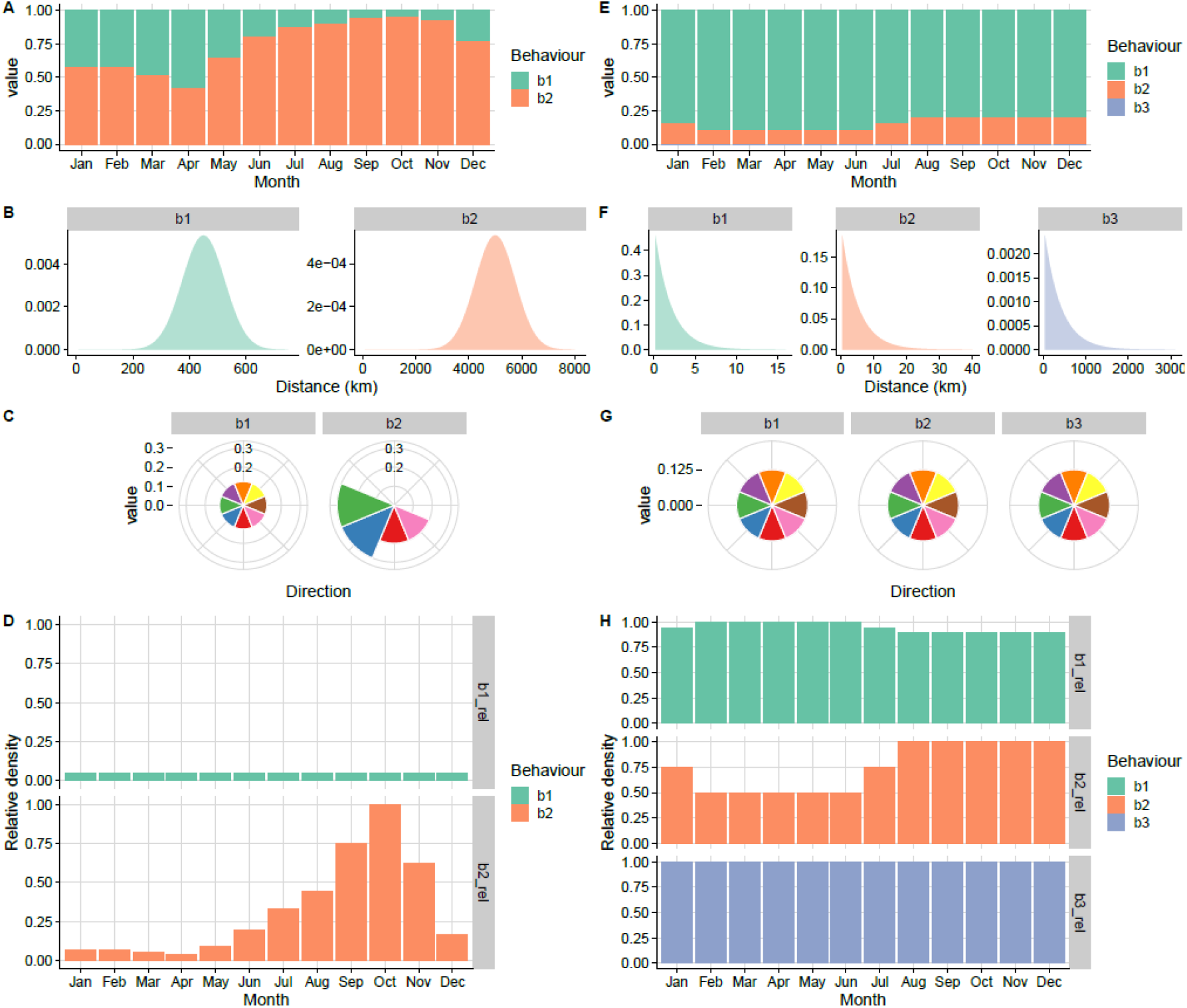
Behavioural parameterisation for highly pathogenic avian influenza (A,B,C,D) and African swine fever (E,F,G,H). A+E. Relative probability between each behaviour group by month (*B*_*abundance*_) (The probability for b3 for African swine fever is 0.001, and thus not visible in this panel). **B+F**. Effective distance (*distance*) travelled by the infection (a combination of animal movement distance and persistence of transmission chains). **C+G**. Probability that an animal or infection ‘moves’ from the outbreak location towards the Netherlands using this cardinal direction (*direction*). The top slice (orange) corresponds with the direction ‘North’; colours provide distinction between the directions. **D+G**. Temporal abundance within behaviour groups (*T*_*abundance*_). Highly pathogenic avian influenza (HPAI); swine fever (ASF); behaviour group (b).

#### African swine fever

To model the entry risk of ASF through wild boar, we defined three behaviour groups: 1) short range movement, 2) young animal dispersal movement, and 3) ‘long jumps’. We defined the latter as outbreaks that occurred more than 250 km from a previous outbreak. For all behaviour groups we assumed that their *directionality* is uniformly distributed, i.e. the probability that the behaviour is in one of eight cardinal or intercardinal directions is 0.125. The short range movement is most likely, with an average probability of 0.85 (between behaviour groups probability based on *B*_*abundance*_, Figure 2E). The young animal movement is less likely, with an average probability of 0.149, and a seasonal influence (Simons et al., 2019). Based on expert input, the peak of this behaviour is when young animals disperse, 1-1.5 year after birth. However, the exact moment at which this occurs varies between years, dependent on factors such as availability of resources, which is reflected in the model by the distribution over time (Figure 2E & 2G). The long jumps are rare (constant probability of 0.001, based on historic observations, see Supplement 2). We parameterised the distance travelled as exponential distributions with parameters that are 1/mean distance of 2, 5, and 388 km, for the three behaviour groups (Figure 2F). The first two were based on the diameter of the home ranges of wild boar (Jerina, Pokorny, & Stergar, 2014; Morelle et al., 2015; Truvé & Lemel, 2003), whereas the distance of the long jumps was based on historical data of ASF in Europe and Russia (Supplement 2). We considered the long jumps as wildlife movements although they are likely to be human-mediated (Guberti, Khomenko, Masiulis, & Kerba, 2019). The temporal abundance (T_abundance_) of wild boar within the behaviour groups is fairly stable over the year, and only affected by the temporal fluctuation of the number of young dispersing animals as described above (Figure 2G). We assumed that all behaviours would pose a risk over the period of 10 days (duration) (Taylor et al., 2020).

### Model assumptions

In the model, we made several assumptions. We considered each outbreak to be of the same size (size = 1), due to a large proportion of outbreaks with missing information on the number of infected animals. This means that now the number of animals is equal to the number of outbreaks, and the overall risk (*R_overall* _*t*_) could also be interpreted as the number of outbreaks reaching the border per time unit. In the baseline model, we made no distinction between outbreaks in ‘wild’ and ‘domestic’ animals, nor did we distinguish between different (bird) species or virus strains. We considered that domestic occurrences of HPAI can be a result of undetected circulation in wild birds, or pose a risk for spill over to wild birds again and as such are also an indication of the infection pressure in wild birds. Similarly, the occurrence of ASF in domestic pigs is likely to be a result from wild boar cases (detected or undetected). Outbreaks are assumed to be representative of disease occurrence, despite known data issues such as heterogeneity in detection/reporting efforts and quality, resulting in underreporting, and bias.

By assuming travel in a cardinal or intercardinal direction, we assumed that travel occurred in a straight line and we did not consider any barriers on the path. Similarly, the distance only influenced whether a case can ‘reach the border’ or not; it did not, for example, affect the probability of introduction success, for example, we did not assume the probability of contact with susceptible animals in the Netherlands.

### Validation

For HPAI in the period 2014-2021, outbreaks in the Netherlands have been observed. Thus, comparison of our model-predicted entry risk with all the observed outbreaks (both in wild and domestic birds in the Netherlands) allowed us to validate the model predictions. We calculated the root-mean-square error (RMSE) to quantify the discrepancy between observed and predicted values. Additionally, we challenged the effect of one of the assumptions we made: restricting the prediction to reported outbreaks that have been marked as ‘wild’ (model 1), instead of considering all reported outbreaks (baseline model).

Validation of results for ASF was not feasible, as no ASF outbreaks were observed in the Netherlands in the period 2014-2021. We therefore also assessed the entry risk of ASF for Belgium and Germany, two countries in which outbreaks did occur. In Belgium the ASF outbreak started in September 2018 (Dellicour et al., 2020); in Germany the first case was detected in September 2020 (Sauter-Louis, Forth, et al., 2021). We ran the model with the same parameters as described above, assuming that the wild boar behaviour did not differ between countries. Distance and direction to the border from the geolocated outbreaks was calculated for Belgium and Germany.

## Results

### Highly pathogenic avian influenza

We assessed the risk of HPAI entry into the Netherlands for the 17,914 HPAI outbreaks that were reported in EMPRES-I worldwide in both wildlife and domestic animals between January 1, 2014 and December 31, 2021. We found that the modelled HPAI entry risk strongly varied between the years (Figure 3); for 2021, we found that a median of 273 outbreaks (*R_overall* _*t*_, 95% uncertainty interval (UI): 254-290) was expected to reach the Dutch border by wild bird migration, for 2019 this was only the case for 1.3 outbreaks (95% UI: 0.6-1.8). The majority of entry risk was due to the long range migration behaviour (b2, 97.8%); the short range flights (b1) contributed to the remaining 2.2% of the risk. The ten countries that contributed most to the risk (ranked by contribution), were Germany, Russia, Denmark, The United Kingdom, Poland, Sweden, Finland, Estonia, Kazakhstan, and the Czech Republic; Ninety-seven percent of the total entry risk came from these ten countries (Supplement 3).

**Figure 3.**
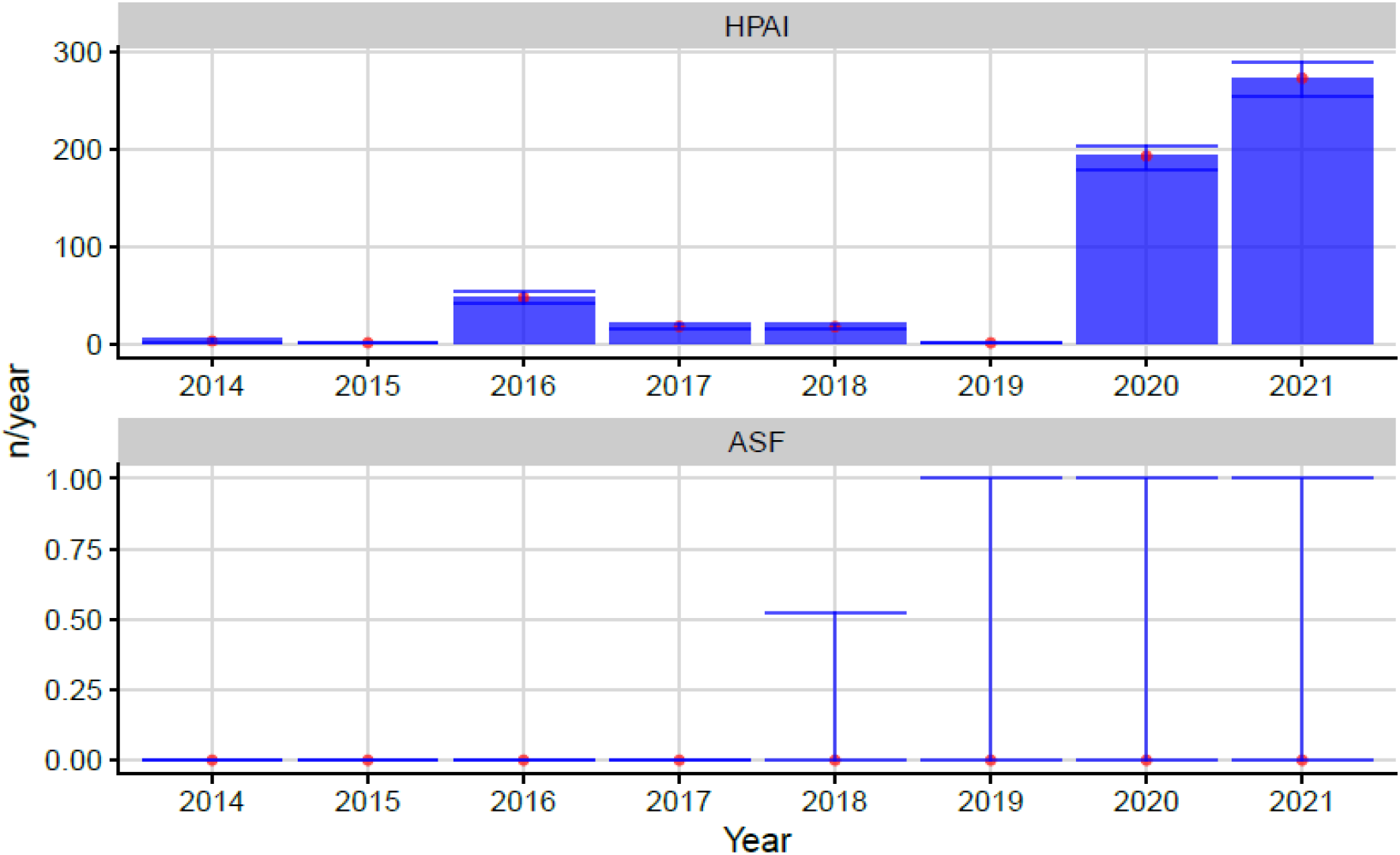
Median yearly risk of entry (, or the number of outbreaks (n) that reach the border per year) of highly pathogenic avian influenza (HPAI) and African swine fever (ASF) through wildlife (bar and point). The error bar provides the 95% uncertainty interval.

### African swine fever

We assessed the entry risk for the Netherlands of 31,451 ASF outbreaks that occurred between January 1, 2014 and December 31, 2021. The yearly risk for ASF entry was low (Figure 3). The median risk was zero for all years, however, the upper 95% uncertainty interval was 1 for 2018 to 2021. All of the entry risk was a result of the ‘long jumps’ (b3).

### Validation

#### HPAI: observed versus predicted

When we compare the estimated yearly risk against the observed number of outbreaks in the Netherlands, for the period 2014-2021 (Figure 4), we find the same pattern. Based on RMSE, the model in which we only considered outbreaks reported in wildlife (model 1; RMSE of 30.1), outperformed the baseline model (model 0; RMSE of 59.5), in which we considered all reported outbreaks (Table 1). However, the alternative model (model 1) underestimated the risk for the years 2014-2018. The improved performance of this model was based on the prediction for the years 2020 and 2021, where the baseline model showed a larger discrepancy between observed and predicted.

**Table 1.** Comparison of different models.

| Model name | Included data/assumptions | RMSE |
| --- | --- | --- |
| Observed | Observed number of outbreaks (wild + non-wildlife) in the Netherlands | Ref. |
| Model 0 (baseline) | Wildlife + non-wildlife outbreaks | 59.5 |
| Model 1 | Wildlife outbreaks only | 30.1 |

**ASF: Comparison The Netherlands, Germany and Belgium.** The entry risk for ASF in Germany started to increase from 2019 onwards, from 0.15 (0.0-2.0) in 2019 to 7.0 (2.0-13.0) in 2021 (Figure 5); In Belgium the modelled risk was low throughout the whole period. When we compared the model results between The Netherlands, Germany and Belgium, we saw that the outbreaks in Germany that started late 2020, were preceded by a period of increased risk; the 2018-2019 outbreak in Belgium did not correlate with the predicted risk. During this period, no outbreaks were observed within the Netherlands.

**Figure 4.**
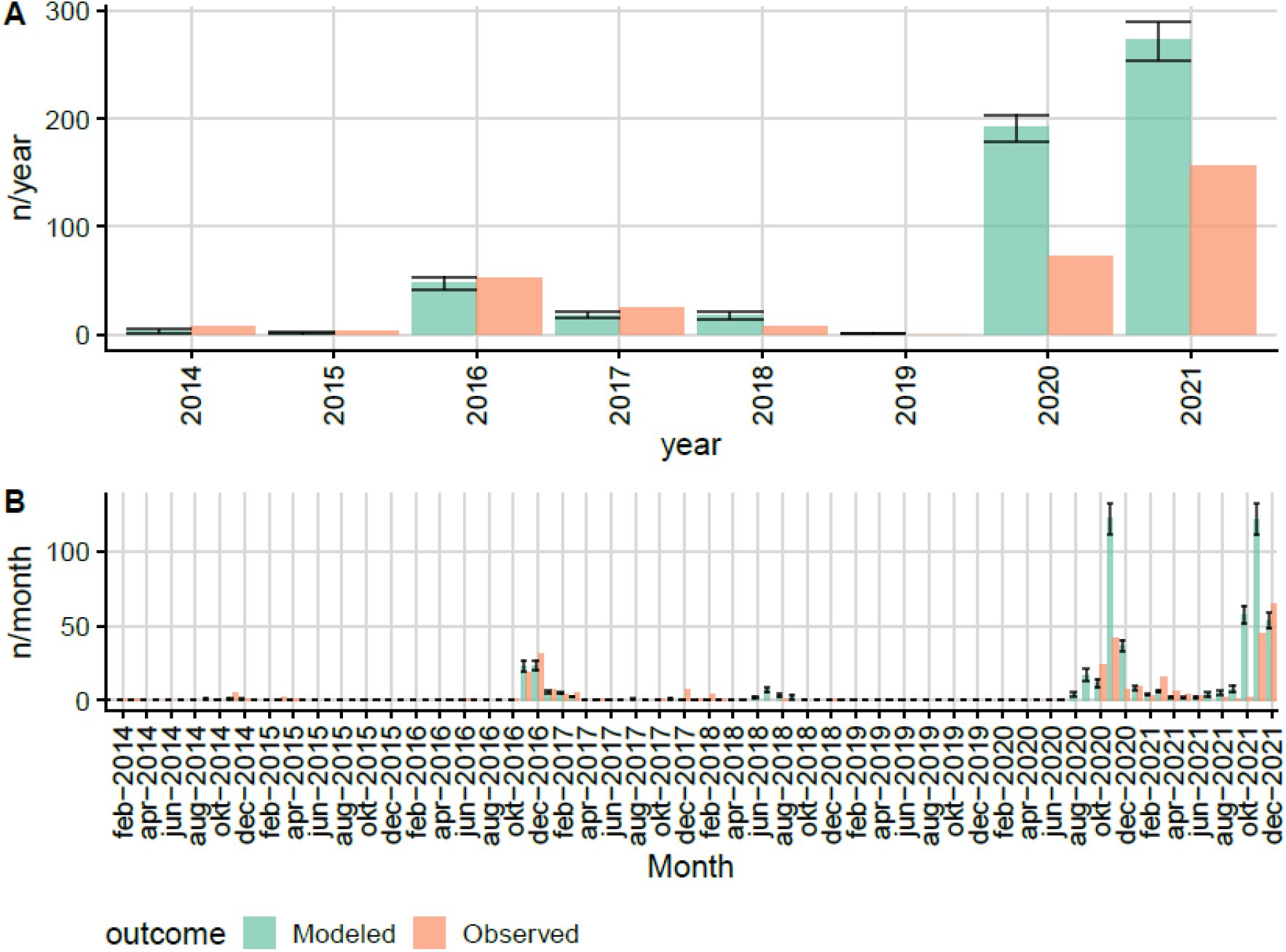
Modelled and observed outbreak risk () for Highly Pathogenic Avian Influenza between 2014-2021. The observed outbreaks are the (wild and domestic) number of outbreaks (n) that were reported in the Netherlands during that time period, by year (A) and by month (B).

#### ASF: Comparison The Netherlands, Germany and Belgium

The entry risk for ASF in Germany started to increase from 2019 onwards, from 0.15 (0.0-2.0) in 2019 to 7.0 (2.0-13.0) in 2021 (Figure 5); In Belgium the modelled risk was low throughout the whole period. When we compared the model results between The Netherlands, Germany and Belgium, we saw that the outbreaks in Germany that started late 2020, were preceded by a period of increased risk; the 2018-2019 outbreak in Belgium did not correlate with the predicted risk. During this period, no outbreaks were observed within the Netherlands.

**Figure 5.**
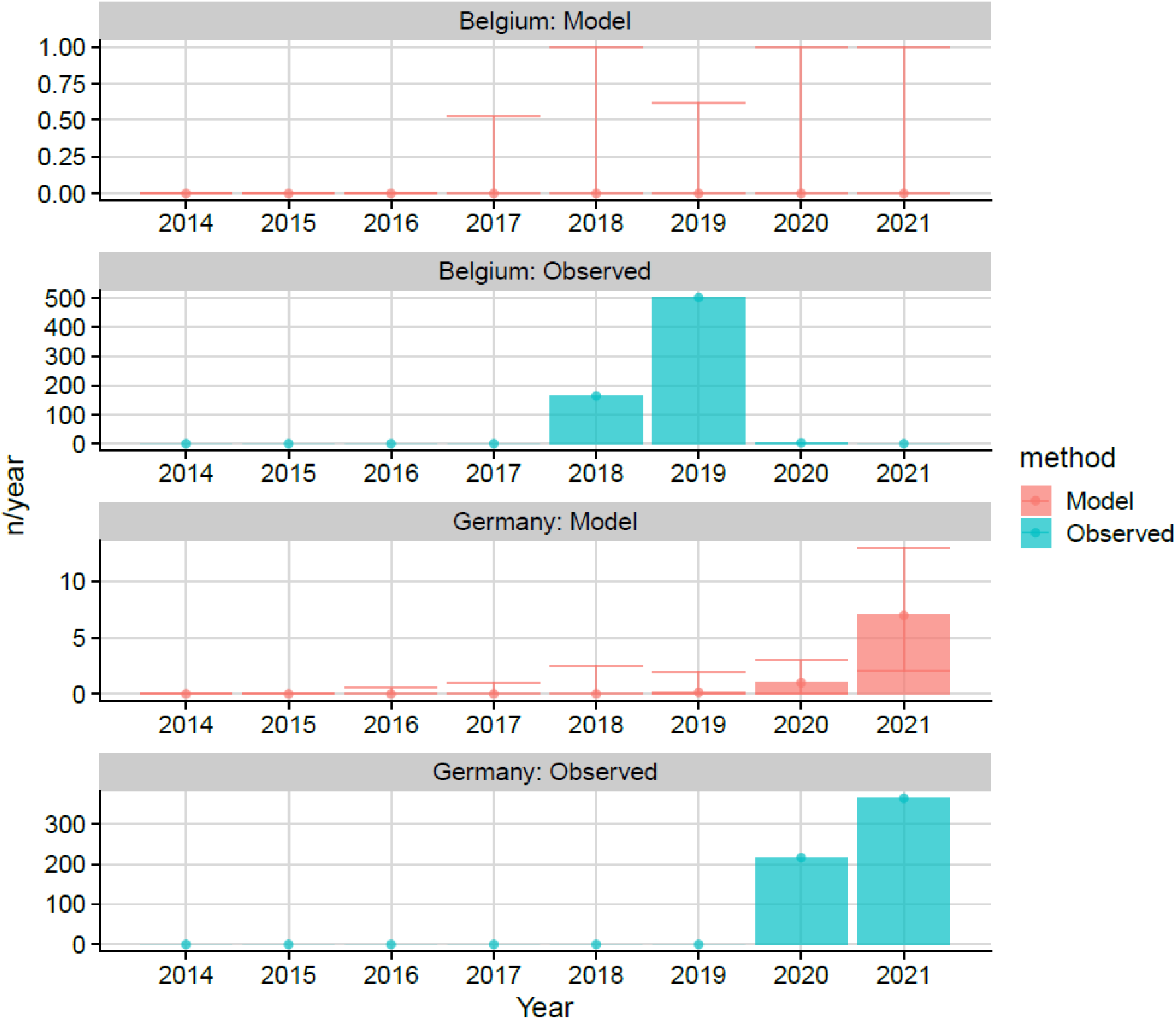
The modelled median yearly entry risk for African swine fever in Belgium and Germany compared to the observed number of outbreaks (n). The error bar provides the 95% uncertainty interval of the modelled risk.

## Discussion

### Summary of findings

Here, we presented a generic entry risk assessment model that provides a flexible and generic solution to estimate the entry risk of animal disease through wildlife movements. We demonstrated that the complexities of animal movements can be reduced to a set of parameters that can be parameterized to mimic animal ‘behaviour groups’. With this, we managed to reach a level of complexity that was necessary to capture the key patterns of wildlife movement, while maintaining flexibility and transparency.

To demonstrate the application of the model, we assessed the entry risk of HPAI through wild birds, and ASF through wild boar for the Netherlands between 2014 and 2021. We found that the risk of HPAI fluctuated between the years with a peak in 2021. The risk of ASF was very low throughout the whole period; only the upper confidence bound indicated a small entry risk. On a yearly scale, the predicted entry risk for HPAI correlated well with observed outbreaks in the Netherlands.

### Interpretation

The difference in risk between HPAI and ASF is driven by the probability that animals reach the Dutch border. Wild boar from ASF outbreaks that occurred between 2014 and 2021 were too far away to pose a risk and the probability of ‘long jumps’ that are most likely the result of human interference was very low; wild birds, through long distance migration, were more likely to reach the border and thus introduce HPAI. This is in line with the recent findings from Engelsma et al. (2022) that describe HPAI outbreaks in the Netherlands as separate introductions originating from wild birds (Engelsma et al., 2022).

If we consider the entry risk for ASF into Belgium and Germany, we found that for Germany the proximity of outbreaks in Poland close to the border caused an increased entry risk before outbreaks were indeed observed. For Belgium, our model did not indicate an elevated entry risk in the time period that the outbreaks in wild boar in Belgium occurred. Previous analyses of these outbreaks confirm that the outbreaks in Germany were likely a result of multiple entries of infected wild boar, whereas the outbreak in Belgium originated from a point-source introduction (Sauter-Louis, Schulz, et al., 2021). The latter is harder to predict due to the stochastic nature of the event.

We should interpret the results semi-quantitatively: The model provides an indication of the relative entry risk over different years rather than a prediction of the absolute number of entries. We should also put the results in perspective of the limitations of the data used. We departed from observed outbreaks of disease, where heterogeneous underreporting and changes in what is reported in time and space is likely. We lack prevalence data, which would allow a more reliable quantitative estimation.

### Strengths and limitations

Our approach has several strengths. First, the model is generic and transparent. We managed to capture the dispersal patterns of HPAI and ASF by wildlife in the same model, despite differences in disease and susceptible species. Second, the model is balanced in its complexity, yet captures the most important patterns that seem to have driven the entry of previous outbreaks. Thus it allowed us to formalize these patterns without the need for a complex model for which data for parametrisation are lacking. Our model has several limitations as well. We had to make several assumptions that have resulted in an oversimplification of the reality. These assumptions might hold true in the context we applied the model, but this might not be the case in other settings. For example, if migration movement in reality is not predominantly in a single direction, the model, by design, failed to capture that pattern. Also, the current parametrisation of HPAI might overestimate the effect outbreaks in neighbouring countries have. Migrating birds are now considered to depart from any location within the distance and direction specified. This means, for example, that autumn occurrence of outbreaks in Germany pose a large risk, where these outbreaks might actually be at end points of migration routes; whereas our model considers these as departure points. Some of the infections in neighbouring countries might even have originated from the Netherlands, rather than posing a risk for entry. This could explain why we saw an increase in the modelled number of infected birds that reach the border in 2020 and 2021, as in recent years, the local circulation of HPAI in Europe has increased, causing the model to predict an increased risk originating from neighbouring countries. Similarly, some of the observed outbreaks in the Netherlands will have resulted from local circulation of the virus rather than new entries. This also demonstrates the limitation of the definition of ‘outbreak’ as reported in the EMPRES-I dataset; many of these events are likely a continuation or re-emergence of an existing event, but the reported outbreaks are not classified as ‘primary’ and ‘secondary’ outbreaks.

Indeed, if we look at the countries that the model predicted were most contributing to the entry risk of HPAI, some might not be in line with the assessment of the phylogenetic trees. Beerens et al (2021) noted that “incursion was not related to viruses detected in eastern Europe, Germany, and Bulgaria earlier in 2020, but was probably associated with fall migration of wild birds to wintering sites in the Netherlands”. Although no HPAI viruses or deaths were reported at wild bird breeding sites in northern Russia, HPAI H5N8 viruses were reported in southern Russia and northern Kazakhstan in September 2020 (Beerens et al., 2021). However, due to selective and limited sequencing of cases, this might also not provide the complete picture of transmission chains.

Endemicity of disease will result in a higher infection risk in the country than accounted for by our model, since the model, as it is parametrized here, only takes into account new introductions and not local circulation. Recently, during 2022, outbreaks of HPAI have occurred outside of the bird migration season, indicating local establishment of the disease in many European countries, including the Netherlands (European Food Safety Authority et al., 2022). Due to the endemicity of HPAI, many of the observed outbreaks are hypothesized to be a result of local circulation rather than new introductions. These changed dynamics, will result in an underestimation of the risk by the current model. However, the model might still be able to predict the number of introductions, although these become less relevant when local circulation is abundant.

### Comparison with other work

There are a few generic quantitative risk models that include wildlife as a pathway for disease entry. Simons et al. (2019) applied their generic risk model to infer the entry risk of ASF, classical swine fever, and rabies through wildlife. They modelled dispersal behaviour of wildlife over a spatial grid, where habitat suitability drives the direction of dispersion. This requires that reliable wildlife density maps are available, which is not always the case. Contrary to our model, they did not apply the model to disease transmitted by birds. However, they do mention that a future extension is possible to include wild birds and also model avian influenza. Taylor et al. (2020) applied a similar approach to Simons et al. (2019), where movement of wild boar depends on habitat suitability. The approaches of Simons et al. (2019) and Taylor et al. (2020) are more demanding than ours with respect to spatial data on susceptible wildlife host populations, which have to be collected for each species separately. The generic framework developed by Taylor et al. (2019) can provide results at different spatial scales, varying from animal holding to country (Taylor, Berriman, Gale, Kelly, & Snary, 2019). Our approach is on the country level, but could relatively easily be adapted to other regional levels as long as file shapes are available. Although we used a quantitative approach to estimate the entry risk by wildlife, results should be used for prioritisation rather than for prediction, similar to some of the semi-quantitative generic tools that have been developed in recent years (Condoleo et al., 2021; Roberts et al., 2011). By calculating the entry risk over multiple years, trends in risk can be observed allowing for horizon scanning and early warning. Our model has been designed such that it can be easily updated when new disease outbreak data become available. It was originally developed as an extension of RRAT, a rapid risk assessment tool to assess the incursion risk of emerging and re-emerging diseases for the Netherlands (de Vos, Petie, van Klink, & Swanenburg, 2022). RRAT only addresses the incursion risk related to human activity, including legal trade in animals and animal products, and animal products illegally carried by travellers, whereas for diseases as ASF and HPAI the incursion risk by wildlife might be more important. A next step would be to merge this tool with RRAT and apply both tools for the same diseases to enable comparisons of the entry risk across pathways and diseases.

### Future improvements

Since models are as good as the data that is put into them, many of the improvements lie in the realm of improving input data. We need to improve the understanding of migration and dispersal patterns. For example, now, migration patterns are based on ringing of birds and retrieval of rings, which is very much dependent on the number and quality of observations, where heterogeneity and underreporting is likely. This results in biased data. Bias that is likely propagated in the models that rely on this data. Translating reliable migration data into probabilities of birds’ departure and arrival would facilitate risk estimation. Additionally, considering the role of landscape and appropriate foraging and resting sites for birds could help to reveal more nuanced migration patterns as well as specific areas at risk. Similarly, extending the model to include barriers such as roads and rivers, and allowing travel across non-linear paths, would refine the prediction of predominantly land-based movement. The framework allows for an extension of the number of behaviours considered, thus enabling the entry risk to be estimated in more detail, for example by bird species or by disease strain or subtype. For example, a more detailed parametrization of different HPAI strains (as different diseases) and different bird species (as different behaviour groups) could increase the accuracy of the model, but does require more detailed outbreak and transmission data. We also need to improve the understanding of disease occurrence. The notification and reporting of disease occurrence is sensitive to differences in reporting across time and space. The probability of a case being detected and reported differs between regions. For ASF, e.g., we see large areas where reporting is absent. Estimation of prevalence of disease would improve the estimation of risk.

A logical extension of the model, is to predict beyond entry. Here, we have only considered outbreaks until the point they reach the Dutch border. Extending the model with the probability of transmission and establishment will give policymakers more insight in the spatial and temporal risk, but requires a better understanding of the movement and behaviour of animals once they have crossed the border. Similarly, the transmission success from wildlife to livestock will depend on their proximity and interaction. For HPAI, extending the model to predict local outbreaks will allow a more reliable comparison with the observed outbreaks as well.

## Conclusion

Movement of wildlife is complex and based on many external factors that vary between regions and time. To assess the entry risk of diseases by wildlife, we described animal movement patterns by a limited number of variables. Each set of parameterized variables represents a ‘behaviour group’, and more nuance and complexity can be added to the model by defining additional ‘behaviour groups’ when more data becomes available. With this approach, we were able to build a generic and flexible framework to assess the entry risk of diseases by wildlife that could be used for both terrestrial animals (illustrated by ASF) and birds (illustrated by HPAI). The model thus enables rapid and transparent estimation of the entry risk for diverse diseases and wildlife species.

## Data availability

All data is available online [https://doi.org/10.7910/DVN/LZUJKS].

## Acknowledgements

We thank Roy Slaterus (Sovon Dutch Centre for Field Ornithology); Armin Elbers, Jose Gonzales, and Nancy Beerens (Wageningen Bioveterinary Research, Wageningen University & Research); Jolianne Rijks (Dutch Wildlife Health Centre); and Dennis Lammertsma, (Wageningen Environmental Research, Wageningen University & Research) for help in estimating input parameters.

This study was funded by the Dutch Ministry of Agriculture, Nature and Food Quality (KB-37-003-033 and WOT-01-003-094).

## Ethical Statement

The authors confirm that the ethical policies of the journal, as noted on the journal’s author guidelines page, have been adhered to. No ethical approval was required as this is a review article with no original research data.

## Author contributions

**MJC:** Conceptualization, Methodology, Modelling, Data Curation, Writing- Original draft preparation **RP**: Interpretation, Data Curation, Visualization, Writing- Reviewing and Editing. **EGMK**: Investigation, Validation, Interpretation. **CV**: Conceptualization, Methodology, Interpretation, Writing- Reviewing and Editing.

