## Supplement for "A generic risk assessment model for animal disease entry through wildlife: The example of highly pathogenic avian influenza and African swine fever in The Netherlands"

### Supplementary information

Supplement 1. Bird migration and parametrisation of highly pathogenic avian influenza

Supplement 2. Long jumps African swine fever

Supplement 3. Contribution by country

#### Supplement 1. Bird migration and parametrisation of highly pathogenic avian influenza

We identified 13 relevant bird species of the Anatidae family, that were used before in the Migration Mapping Tool (<https://app.bto.org/ai-eu/main/data-home.jsp>) with the exception of the Brant Goose. We selected the Greater White-fronted Goose (*Anser albifrons),* Eurasian Wigeon (*Anas penelope*), Gadwall (*Anas strepera*), Common Teal (*Anas crecca*), Mallard (*Anas platyrhynchos*), Northern Pintail (*Anas acuta*), Garganey (*Anas querquedula*), Northern Shoveler (*Anas clypeata*), Red-crested Pochard (*Netta rufina*), Common Pochard (*Aythya ferina*), Tufted Duck (*Aythya fuligula*), Common Coot (*Fulica atra*), and the Brant Goose (*Branta bernicla*). These species are both abundant in the Netherlands and can act as potential host for HPAI. Most species have their wintering grounds in the Netherlands, except for the Garganey (Figure S1.1). In general, their breeding grounds are located North – East of the Netherlands (see below: ‘The origin of the birds’).

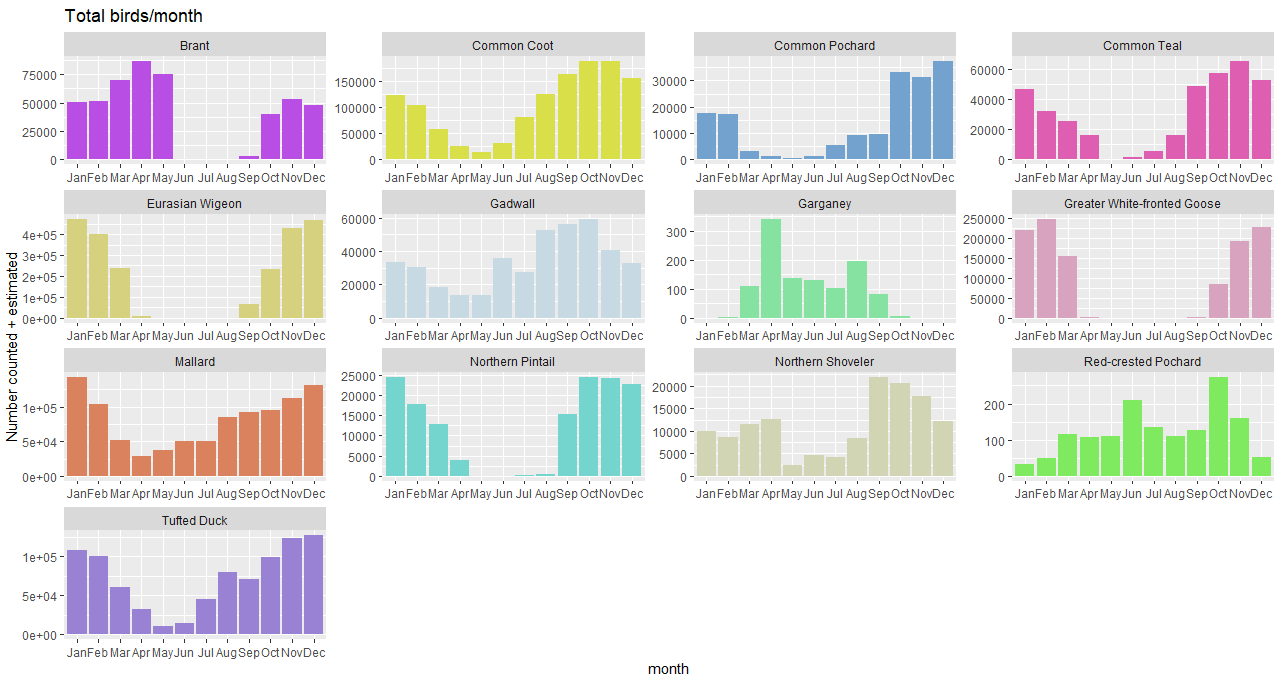

**Figure S1.1.** The monthly number of counted and estimated birds by species. Source: SOVON (https://stats.sovon.nl/).

##### Data processing

The relevant period for the risk of incursion is when the number of animals of a certain wild bird species in the Netherlands increases. Of the 13 identified species, we retrieved the estimated number of animals per month as provided by ‘Sovon Vogelonderzoek Nederland’ (<https://stats.sovon.nl/>). We calculated the ‘influx’ of animals as the difference in the number of animals in the current and the previous month. Since only the influx of animals means incursion risk, we censored the calculated influx on positive values by species. Then, we aggregated the positive influx of all species per month. The relative influx per month is calculated as a proportion of the maximum influx which happened to be in October (figure, table). We considered that, on average, the birds depart 15 days earlier from their country of origin than that they arrive in the Netherlands. We use this ‘influx shift’ since disease is observed in the country of origin.

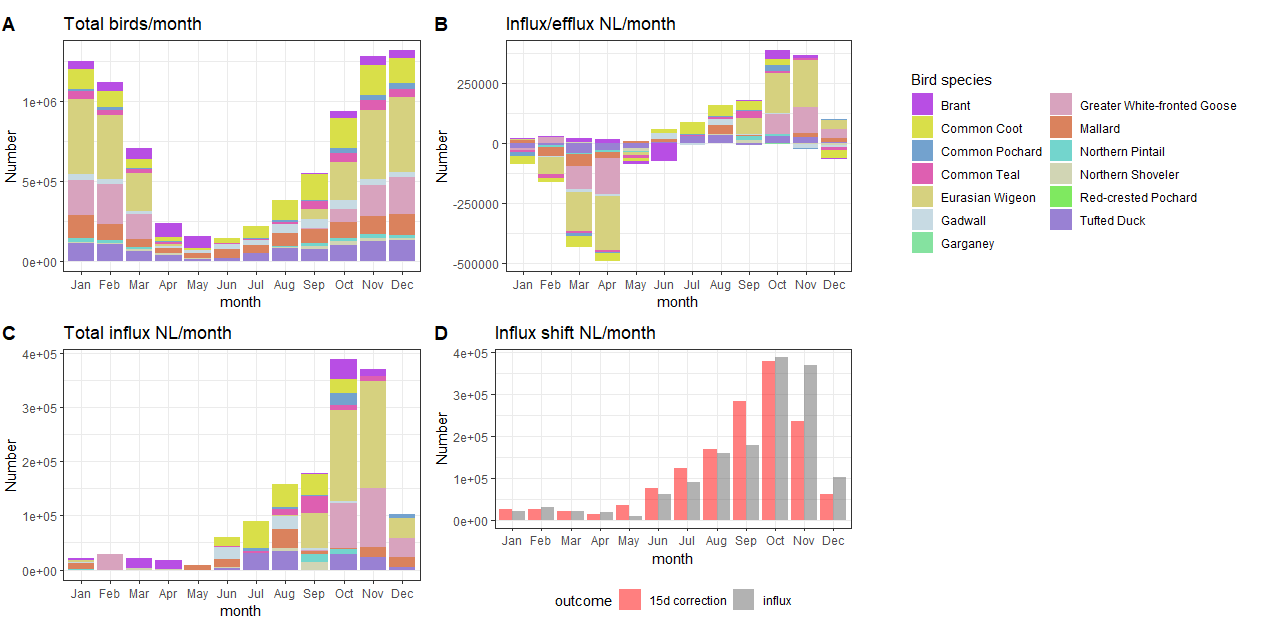

**Figure S1.2.** The monthly number of counted and estimated birds by species in the Netherlands. A. Cumulative number of birds by month; B. The difference in number of birds between month (influx and efflux). C. The positive difference or influx of total birds of the 13 selected species. D. The influx of the 13 selected species combined, shifted 15 days earlier (red bars) to reflect departure from country of origin, the grey bars represent the total influx from panel C. Data source: SOVON (https://stats.sovon.nl/).

##### The origin of the birds

A commonly used method to track migration patterns of birds is ringing birds and observing or capturing these birds at a later time. These observations provide some insights in where birds travel. However, the probability of observations is not homogeneous and thus patterns suffer from bias. We explored the migration patterns as described in the Migration Mapping tool (<https://euring.org/research/migration-mapping-tool>). Based on these migrations maps of the 13 selected species, we reduced the long distance movement (migration) of the selected birds to be arriving from a predominantly North, North-East, and East direction.

#### Supplement 2. Long jumps African swine fever

To infer the risk of ‘long jumps’ of African swine fever (ASF), we analysed the outbreaks that occurred over the last decade. Between 30 April, 2010 and 14 September 2021, 24,191 ASF outbreaks were reported on the European continent. We calculated for all outbreaks the distance to all previous outbreaks. Eight of these outbreaks had a minimum distance between previous outbreaks of more than 250 km. This results in a probability of 3.3 per 10,000 outbreaks that a ‘long jump’ occurs. We considered a worst case scenario where we rounded this to a probability of 0.001 (10/10,000). The median distance of these long jumps was 388 km (IQR: 304-521 km) (Figure S1.3).

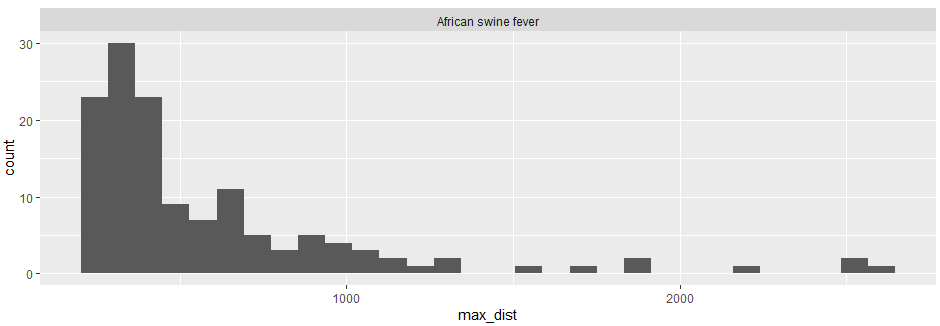

**Figure S1.3** Frequency of outbreaks with a outbreaks had a minimum distance between previous outbreaks of more than 250 km.

#### Supplement 3. Contribution by country

Here we provide the total number of predicted outbreaks (median, and 95% uncertainty interval) that have reached the border of the Netherlands between January 1, 2014 and December 31, 2021.

**Table S3.1.** Median number of highly pathogenic avian influenza outbreaks (success) by country that were predicted to reach the Netherlands between 2014 and 2021 and the 95% uncertainty interval.

| **country** | **Success, median** | **2.5 percentile** | **97.5 percentile** |
| --- | --- | --- | --- |
| Germany | 260.90 | 243.93 | 278.06 |
| Russian Federation | 100.67 | 88.29 | 110.83 |
| Denmark | 65.52 | 57.23 | 77.28 |
| U.K. of Great Britain and Northern Ireland | 42.29 | 30.59 | 50.68 |
| Poland | 27.40 | 23.52 | 30.91 |
| Sweden | 19.09 | 14.04 | 23.02 |
| Finland | 18.25 | 13.87 | 23.49 |
| Estonia | 6.54 | 3.65 | 9.27 |
| Kazakhstan | 6.02 | 3.74 | 9.23 |
| Czech Republic | 3.96 | 1.67 | 5.65 |
| Norway | 3.50 | 1.38 | 6.23 |
| Ukraine | 2.76 | 0.91 | 3.91 |
| Ireland | 2.19 | 0.16 | 4.22 |
| Lithuania | 1.47 | 0.79 | 2.01 |
| Austria | 1.16 | 0.25 | 1.95 |
| China | 0.90 | 0 | 2.66 |
| Faroe Islands | 0.75 | 0 | 1.50 |
| Latvia | 0.46 | 0.07 | 0.86 |
| Belgium | 0.35 | 0 | 1.51 |
| Slovakia | 0.33 | 0.10 | 0.63 |
| France | 0.29 | 0 | 1.69 |
| Romania | 0.16 | 0 | 0.16 |
| Hungary | 0.07 | 0 | 0.13 |
| Switzerland | 0.06 | 0 | 1.56 |

**Table S3.2.** Median number of African swine fever outbreaks (success) by country that was predicted to reach the Netherlands between 2014 and 2021 and the 95% uncertainty interval.

| **country** | **Success, median** | **2.5 percentile** | **97.5 percentile** |
| --- | --- | --- | --- |
| Poland | 0 | 0 | 2 |
| Belgium | 0 | 0 | 1 |
| Estonia | 0 | 0 | 1 |
| Germany | 0 | 0 | 1 |
| Latvia | 0 | 0 | 1 |
| Romania | 0 | 0 | 1 |
